# Backmapping with Mapping and Isomeric Information

**DOI:** 10.1101/2023.09.05.556379

**Authors:** Siyoung Kim

**Affiliations:** Pritzker School of Molecular Engineering, University of Chicago, Chicago, IL 60637 USA

## Abstract

I present a powerful and flexible backmapping tool named Multiscale Simulation Tool (`mstool`) that converts a coarse-grained (CG) system into all-atom (AA) resolution and only requires AA to CG mapping and isomeric information (*cis/trans/dihedral/chiral*). The backmapping procedure includes two simple steps: a) AA atoms are randomly placed near the corresponding CG beads according to the provided mapping scheme. b) Energy minimization is performed with two modifications in the AA force field (FF). First, nonbonded interactions are replaced with cosine functions to ensure numerical stability. Second, additional torsions are imposed to maintain molecules’ isomeric properties. To test the simplicity and robustness of the tool, I backmapped multiple membrane and protein CG structures into AA resolution, including a four-bead CG lipid model (resolution increased by a factor of 34) without using intermediate resolution. The tool is freely available at github.com/ksy141/mstool.

**TABLE OF CONTENTS (TOC):** 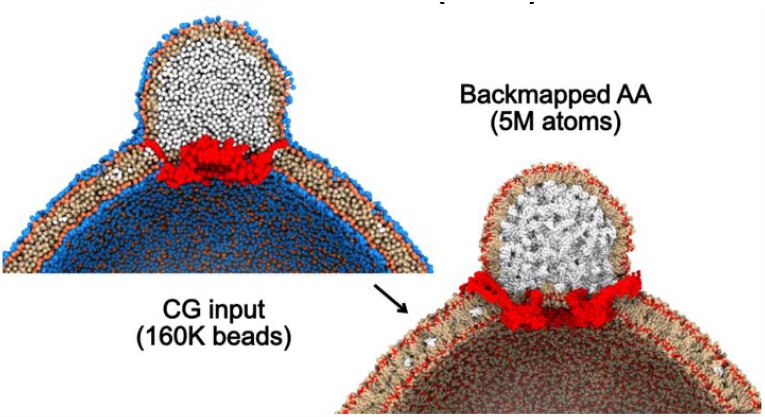

## INTRODUCTION

CG molecular dynamics (MD) simulations have become a popular method in studying biological processes with time and length scales beyond the reach of AA-MD simulations,^1–6^ such as large-scale membrane deformation, organelle biogenesis, and the complete virion or capsid of a virus.^7–10^ In this context, resolution transformation from CG to AA, or backmapping, is necessary to obtain detailed atomistic interactions between molecules, a level of detail not available in CG resolution. Therefore, CG-MD and backmapping can complement each other: one can perform preliminary CG-MD simulations to navigate systems’ equilibrium distribution, followed by additional AA-MD simulations, using initial structures obtained from backmapping.

While it is straightforward to map AA structures into CG ones, the reverse requires recovery of the degree of freedom that has been integrated away.^11,12^ One of the approaches is fragment-based reconstruction, which replaces CG beads with the corresponding atomistic fragment. This popular approach has been used in protein structure modeling by reconstructing full atomistic protein structures from their alpha carbon (CA) positions. For example, a four-residue backbone fragment that fits the four consecutive CA atoms from the Protein Data Bank (PDB) search is used to construct backbone atoms, followed by searching for the most probable side chain conformation.^13,14^ The same approach has been used on protein and DNA complexes.^15^ A more flexible and general fragment-based approach has been developed by Stansfeld and colleagues.^16,17^ This approach can convert CG-MD structures of membrane protein and lipids at Martini resolution^18–21^ to AA structures. Throughout this manuscript, the latest version of their tool will be referred to as `CG2AT2`.^17^

There is another approach in which atoms are positioned based on local geometrical information that users provide. For instance, `CG2AA` uses the positions of three consecutive CA atoms to construct backbone and side chains based on a simple geometrical algorithm.^22^ Wassenaar et al. have defined the relative positions of atoms not only for protein but also for lipids, extending applications of the geometrical approach.^23^ This tool will be referred to as `Backward` in this manuscript. Finally, there are recent studies that use machine learning for resolution transformation.^24–27^

The above approaches have focused on constructing initial AA structures from CG structures, followed by standard energy minimization. This paper presents a backmapping approach that uses a modified FF in energy minimization. It is powerful enough that the initial positions of atoms can be random without fragment alignment or geometrical projection, greatly simplifying user inputs into a minimal set of information: mapping scheme and isomer properties. The tool replaces nonbonded interactions with cosine functions to ensure numerical stability of energy minimization while keeping bonded interactions intact. In other words, a system is relaxed with higher priority given to bonded interactions, while cosine nonbonded interactions ensure no atoms overlap. In addition, a set of torsion potentials is imposed during relaxation to maintain the provided isomeric properties of molecules (*cis/trans/chiral/dihedral*). While the idea is simple, the tool is powerful and capable of constructing complete AA structures from large-scale, highly CG systems.

The rest of the paper is structured as follows. I will first describe a modified FF. In Results, I will test whether torsions applied to the provided isomeric properties work as intended. Then, I will compare the backmapping performance of `mstool` with the popular geometrical projection approach and fragment-based approach, `Backward`^23^ and `CG2AT2`,^17^ respectively. I will then present more examples of resolution transformation of CG lipid and/or protein systems. The examples described in this paper are available at the Github link with a step-by-step guideline.

## METHODS

For each type of molecule (residue) present in CG structures, a mapping scheme describes which AA atoms belong to which CG beads. It also should include molecules’ isomeric properties such as *cis, trans, dihedral*, and/or *chiral*. The predefined mapping files based on the Martini FF are read if no user mapping scheme is provided.^18–21^

The backmapping procedure consists of three steps: Ungrouping, relaxing a system, and checking a backmapped structure. The first step places AA atoms near their corresponding CG beads according to the provided mapping scheme, implemented as `mstool.Ungroup`. The output of this step is an intermediate AA structure, but not at energy minimum because AA atoms are clustered at the locations of CG beads. Water CG beads are treated separately in this step because each water bead represents more than one water molecule in most CG models. For instance, the Martini FF contains four water molecules per each water bead. Users can provide the number of water molecules that are represented by each water CG bead and the residue name of CG water. If no user input is given for water, they are set to 4 and W, respectively, consistent with the Martini FF convention.

The next step is called Reduced Nonbonded Energy Minimization (REM), implemented as `mstool.REM`. A system is energy minimized with a modified FF. First, CHARMM36m protein^28^ and CHARMM36 lipid^29^ FFs are applied to the output structure of `mstool.Ungroup`. Then, the tool replaces the positive part of LJ potential (U > 0) with cosine repulsion, expressed as

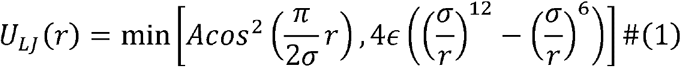

A charged interaction is described by

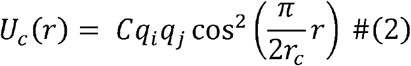

By default, *A =* 100 *kJ/mol, C=* 50 *kJ/mol*. Nonbonded interactions are cut-offed at *r*_*c*_ = 1.2 *nm*. Eqs. 1 and 2 are illustrated in Fig. 1A. The modified nonbonded interactions ensure energy and force values do not diverge even if atoms are very close and smooth potential energy surfaces, making energy minimization easier.

**Figure 1.**
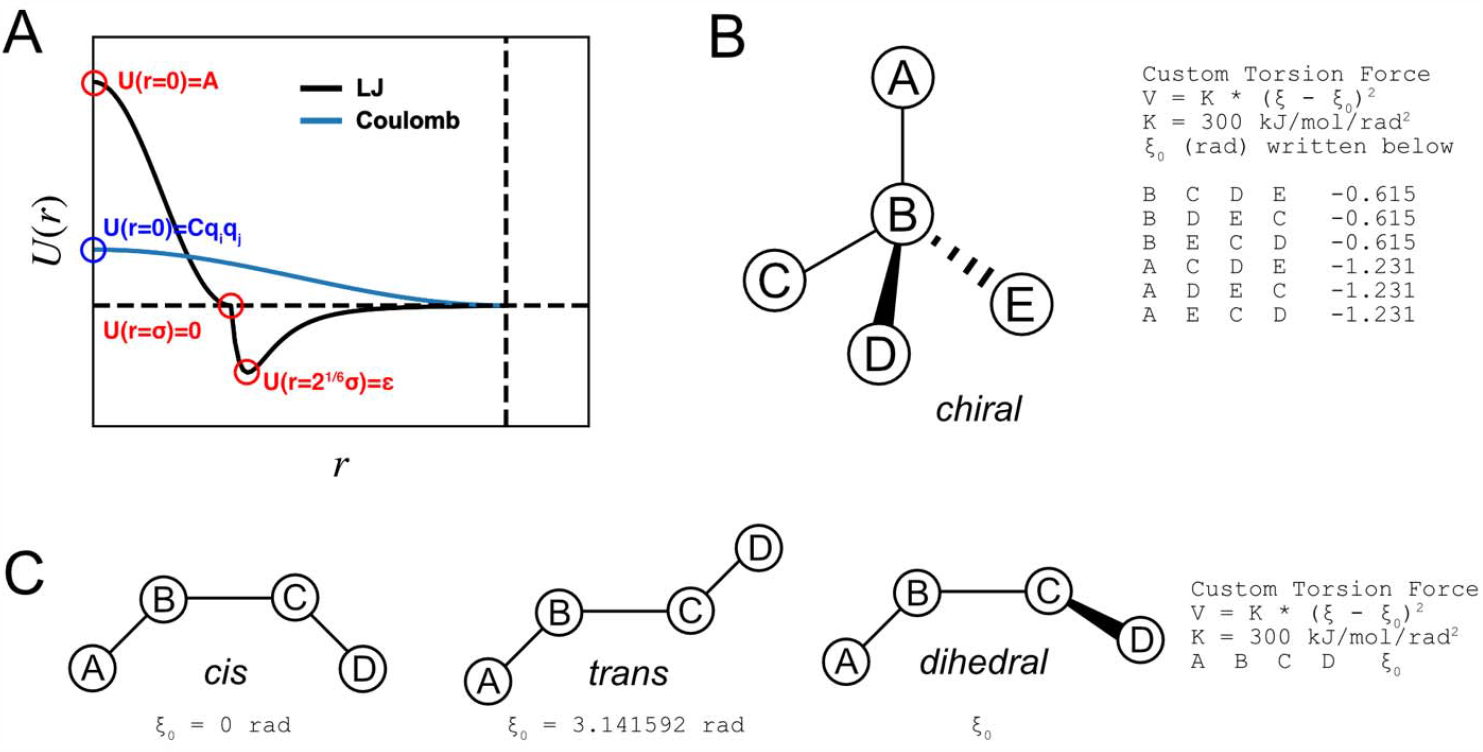
Illustration of the modified force field. (A) Nonbonded interactions. (B) Torsions for *chiral*. (C) Torsions for *cis, trans*, or *dihedral*.

In addition to using modified nonbonded interactions, isomeric torsion terms are added during REM to maintain the provided isomeric properties of molecules, which should be written in mapping files. A set of improper dihedral terms is applied for *chiral* (Fig. 1B). A single improper dihedral term is applied for each *cis, trans*, or *dihedral* (Fig. 1C), described by *V* = *K* (*ξ* − *ξ*_0_)^2^. For *dihedral, ξ*_0_ is determined by user input. For *cis* or *trans, ξ*_0_ = 0° (*cis*) *or* 180° (*trans*). *K* is set to 300 *kJ/mol/rad*^2^.

Every amino acid except for glycine has a *chiral* center at the carbon alpha (CA) atom, and their chirality should be denoted in a mapping scheme. However, a peptide bond, which is dominantly in the *trans* configuration, cannot be specified in a mapping scheme because it is a cross-residue property and cannot be defined within a single residue. To prevent any *cis* peptide bonds, `mstool` applies a *trans* dihedral (Fig. 1C) in every peptide bond that it detects using atomic names during REM. With the modified FF, `mstool` uses openMM for energy minimization.^30,31^ The search of a local minimum is performed with the L-BFGS algorithm until the root-mean-square value of all force components reaches 10 kJ/mol/nm.^32^

The last step in the backmapping procedure is checking the resulting AA structures, implemented as `mstool.CheckStructure`. It reports which isomeric properties are reviewed (written in mapping schemes) and flipped (inconsistent with mapping schemes) isomeric properties. This function only checks the defined isomeric properties in the provided mapping schemes. Unspecified isomeric properties will not be detected or checked. However, `mstool.CheckStructure` automatically detects and checks *cis* peptide bonds because a peptide bond is a cross-residue property and cannot be specified in a mapping file.

When using the `mstool` backmapping, a system should not have two or more residues with the same residue name, residue number, and chain name. This may be an issue for a very large CG system because the residue number is limited to up to only four digits (0-9999), and the chain name can be defined with only one character in a PDB format. To solve this issue, `mstool` also accepts a Desmond structure file (DMS), which has no limits on the length of residue name, residue number, and chain name.^33^ Finally, one CG bead should not represent two or more molecules except water, which is treated separately inside the tool. Therefore, resolution transformation of supra-CG or mesoscale CG models is not supported.

All the backmapping and simulations were performed on a MacBook Pro (16-inch, 2021) with an Apple M1 Pro chip. For NPT simulations, Langevin dynamics was used with a target temperature of 310 K and a friction coefficient of 1 ps^-1^. A semi-isotropic or isotropic Monte Carlo barostat was used with a target pressure of 1 bar and a pressure coupling frequency of 100 steps.^34,35^ A cutoff of 1.2 nm was used for the direct space interactions, and particle mesh Ewald was used for the long-range electrostatic interactions.^36^ All bonds involving hydrogen atoms were constrained.^37^ Simulations were evolved with a 2 fs time step. Simulations and protein dihedrals were analyzed using MDAnalysis.^38,39^ The C36-c parameters were used for triolein.^40^ Structures were visualized with ChimeraX.^41,42^

## RESULTS

### Enantiomers and Configurational Isomers

Due to their coarse nature, CG models contain less isomeric information than their AA counterparts. For example, the CA atom of every amino acid except for glycine is a *chiral* center, and peptide bonds are dominantly in *trans* configuration at AA resolution. However, many CG models, including Martini, have only one CG backbone bead, removing both enantiomeric and configurational isomerism. For this reason, care must be taken when reconstructing AA models from CG ones because the resulting AA structures should have the correct isomeric properties.

In `mstool`, torsions are imposed to ensure backmapped molecules have correct isomeric properties during REM. To test whether these torsions work properly, I backmapped CG systems with and without providing isomeric properties. The first test system was a Martini bilayer membrane with each leaflet constructed from 20 POPC molecules. Each POPC molecule has one *chiral* center and one *cis* configuration (Fig. 2A). The CG and backmapped structures are shown in Fig. 2B.

**Figure 2.**
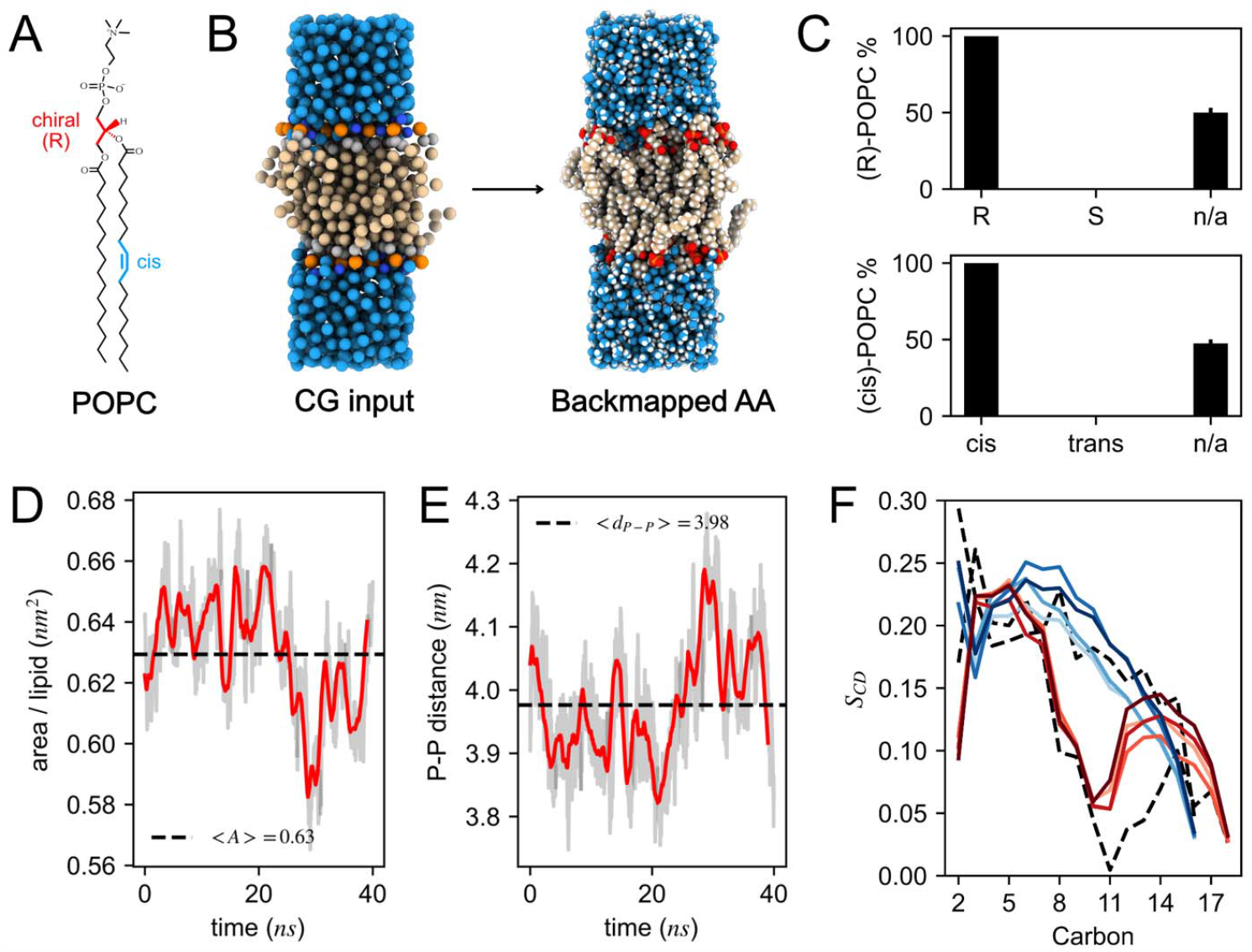
Backmapping and simulation of POPC bilayer membrane. (A) Molecular structure of POPC (B) CG and AA structures (C) Isomeric properties of backmapped POPC molecules (y-axis) with given isomeric properties (x-axis). n/a represents unspecified isomeric properties. Error bars are standard errors of four replicates. (D) Area per lipid. (E) Distance of phosphate across bilayer in AA-MD simulation. Gray lines are the instantaneous values at 20 ps intervals. Red lines are the moving average with 1 ns window. Dashed lines are the averaged values. (F) Order parameters for *sn*-1 (blue) and *sn*-2 (red) chains, depicted every 10 ns, with darker color representing later simulation time. Dashed lines indicate the order parameters of the initial structure.

When the *chiral* center was given in a mapping file, either R or S, all the backmapped POPC molecules were enantiomerically pure with their chirality as provided. If the *chiral* center was unspecified, the tool produced a racemic mixture (Fig. 2C). Similarly, when the geometric isomerism was specified, either *cis* or *trans*, all the molecules had homoconfiguration. When configuration was not specified, an equal mixture of *cis* and *trans* molecules was created (Fig. 2C). The important conclusion from this test is twofold: A) Enantiomeric and configurational isomeric properties must be provided to have desired isomeric properties in resulting AA systems. B) Imposed torsions play a critical role in reconstructing molecules with desired enantiomeric and configurational properties.

### Lipid Properties

I performed 40 ns of NPT simulation of a POPC bilayer membrane, backmapped with the correct enantiomeric (R) and configurational (*cis*) properties, and then analyzed the physical properties of the membrane. The initial values of the area per lipid (Fig. 2D) and distance between phosphate atoms across the bilayer (Fig. 2E) were already in the equilibrium region. This is because the Martini FF agrees with the experimental and AA-MD simulation data.^18–21^ The order parameters, *S*_CD_ = 0.5 × | < 3 *cos*^2^Θ − 1 > |, where Θ is the angle between CH vectors with the bilayer normal, are more directly related to the backmapping performance. In this POPC bilayer membrane example, the initial structure, backmapped from the CG structure, already had the reasonable order parameters and could distinguish *sn*-1 and *sn*-2 chains (dashed lines in Fig. 2F). This is a reinsuring result given that the backmapping procedure does not involve any NPT/NVT simulations but only energy minimization. Furthermore, the first 10 ns simulation could already equilibrate the bilayer membrane (Fig. 2F).

### Comparison with Backward and CG2AT2

The backmapping performance of `mstool` was compared with the two other popular tools, `Backward`^23^ and `CG2AT2`.^17^ The first system was a Martini bilayer membrane of POPC and DOPC of various sizes (Fig. 3A). For each system size, four equilibrated Martini bilayer membranes were prepared and then backmapped with `Backward`, `CG2AT2`, and `mstool`. While all tools can backmap the CG systems into AA structures, `Backward` produced flipped configuration, and `CG2AT2` flipped enantiomers. In contrast, all molecules converted with `mstool` preserved their isomeric properties.

**Figure 3.**
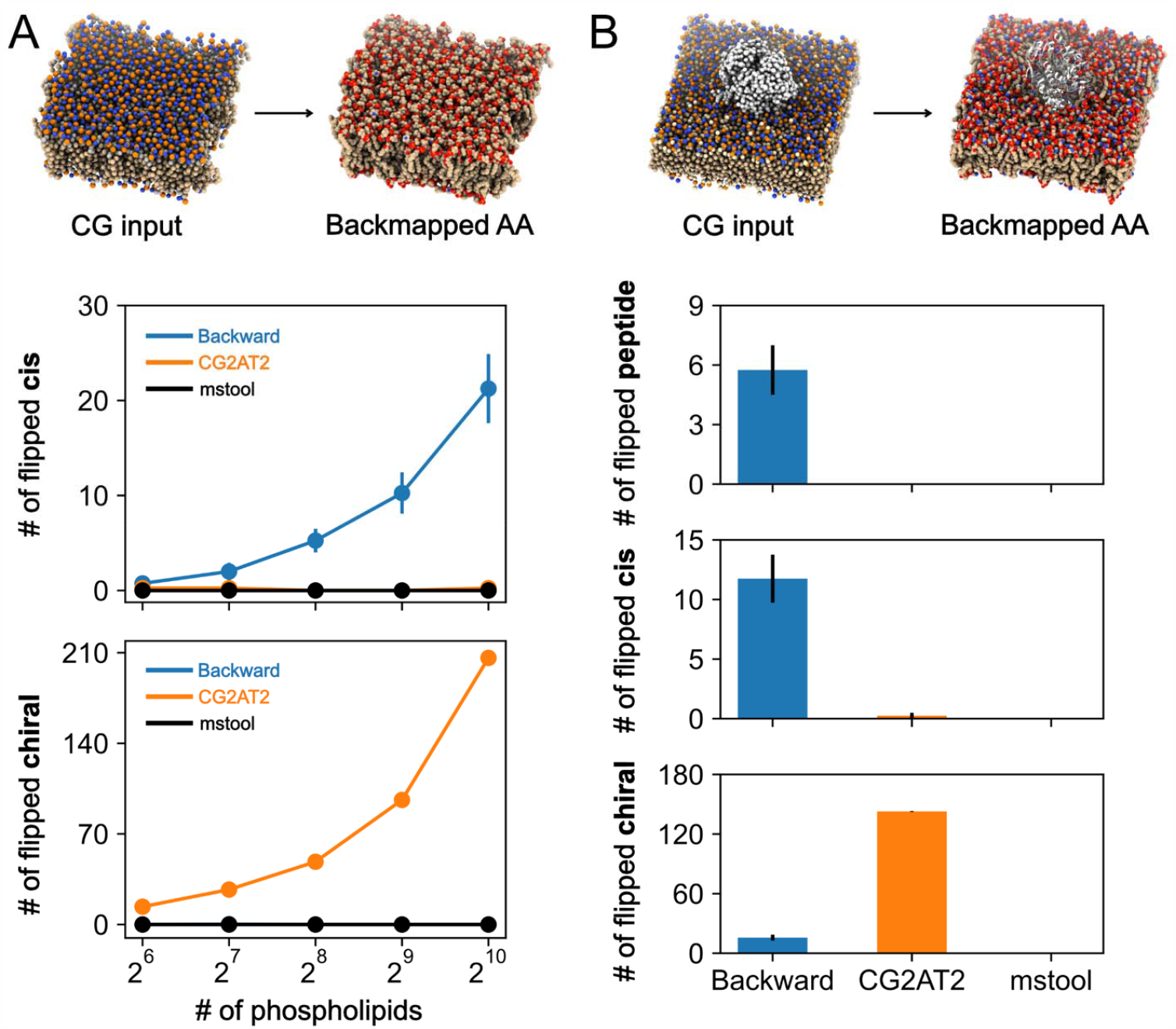
Comparison of backmapping performance. (A) DOPC and POPC bilayer membrane (B) WzmWzt ABC transporter in POPE and POPG membrane. The number of flipped enantiomerical and configurational properties are shown. Peptide on the right means the number of *cis* peptide bonds. Error bars represent standard errors of four backmapping results. Hydrogen atoms in backmapped AA structures are omitted for visual clarity from now on.

The second system was the WzmWzt ABC transporter (PDB: 6M96), a membrane protein system.^44^ Its CG structure was provided by `CG2AT2`, with the protein surrounded by 504 POPE and 125 POPG lipid molecules (Fig. 3B).^17^ The CG structure was backmapped four times using the three tools. Consistent with the previous example, cis-to-trans flipped configurations and flipped enantiomers were observed in AA structures when backmapped with `Backward` and `CG2AT2`, respectively. In the `Backward` structures, *cis* peptide bonds were also detected. There were no flipped isomeric properties for the `mstool` structures.

### Conformation

Unlike enantiomeric and configurational isomerism, conformational isomerism changes during unbiased AA-MD simulations. In fact, the boat-to-chair conversion or the interconversion of twist-boat is frequent. However, neither the ring inversion (chair-to-chair) nor the chair-to-boat conversion is likely probable in unbiased AA-MD simulations because of the stability of the chair state. Therefore, the most probable conformation should be specified using *dihedral* torsions during backmapping.

Cyclohexane was used as a test system to evaluate the backmapping tool’s performance of conserving conformational isomerism. Cyclohexane has multiple conformations: chair (C), boat (B), twist-boat (TB), and half-chair (HC) as shown in Fig. 4A. A total of 125 cyclohexane molecules, each of which was represented by one CG bead, was backmapped with and without *dihedral* torsions and equilibrated for 10 ns in NPT ensemble (Fig. 4B). When *dihedral* torsions were provided, an initial conformation of all cyclohexane molecule was consistent with the provided torsions (Figs. 4C-4E). For instance, if a chair conformation was specified in a mapping file, all the cyclohexane molecules were initiated with the chair conformation (red dots in Figs. 4C and 4D). Given the stability of the chair conformation, all the molecules maintained their chair conformation during the NPT simulation (heatmap in Figs. 4C and 4D). If a twist boat conformation was specified, the initial conformation of cyclohexane was a twist boat. However, during the NPT equilibrium, cyclohexane changed its conformation to the most stable chair conformation, either C1 or C2 (Fig. 4E).

**Figure 4.**
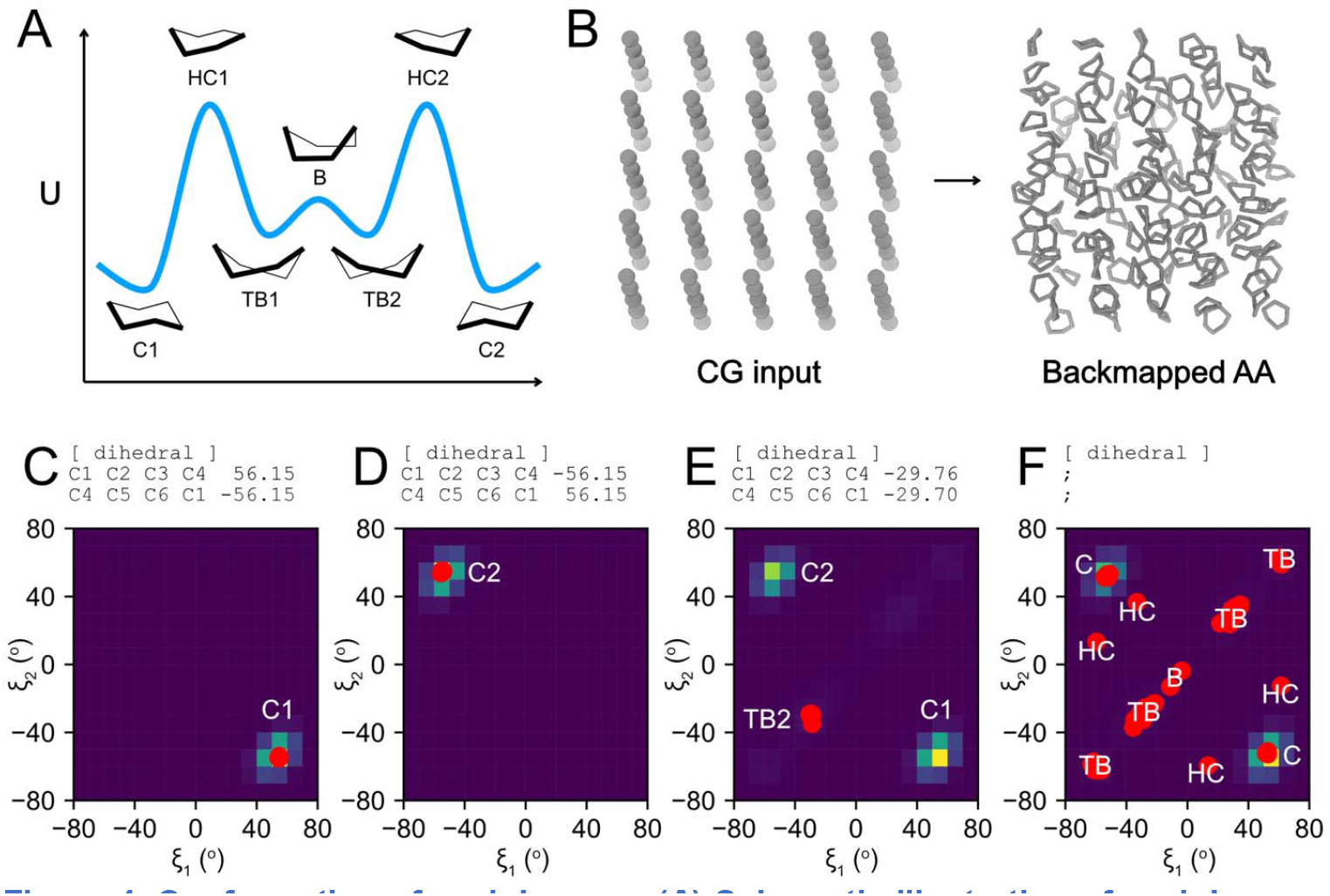
Conformation of cyclohexane. (A) Schematic illustration of cyclohexane conformation energy landscape. (B) CG and AA structures. (C-F) Dihedrals of cyclohexane molecules. Red dots indicate the conformations of backmapped structures. Heatmaps indicate the conformations during the following NPT simulations. Brighter colors represent more populated states in the heatmaps.

When isomeric properties were not given, unlike the previous example that had an equal mixture of *cis/trans* and *R/S* (Fig. 2C), the conformations of cyclohexane were not equally distributed: twist-boat was 80.8%, chair 14.4%, half-chair 3.2%, and boat 1.6% (Fig. 4F). The chair conformation is the most stable state and should be the most probable state. However, the twist boat conformation was the most frequent conformation when torsions were not provided, suggesting that the resulting structure was not Boltzmann distributed.

There are multiple reasons for the non-Boltzmann distribution of conformations. First, the 1-4 repulsion is a large energetic component that makes the chair conformation lower energy than the boat conformation. In the `mstool` pipeline, the repulsion in the modified FF is much softer than the original LJ. Second, backmapping involves energy minimization but no dynamics. Therefore, no information on temperature is given during REM. Therefore, the softer repulsion in the modified FF combined with the lack of dynamics results in the non-Boltzmann distribution of conformations.

However, during the following NPT simulation, all the conformations changed to the chair conformation (C1 or C2), as shown in the heatmap of Fig. 4F. Therefore, it should be noted that backmapping does not produce Boltzmann distributed initial conformations and should be only used to make an initial structure for AA-MD simulations which then can be further equilibrated using NPT or NVT simulation.

Finally, it should be noted that cyclohexane is rather a simple molecule, and its chair conformations are all equivalent. Other ring molecules (e.g., glycosylated residues or inositol-containing lipids) have multiple chair conformations with different energetic levels. Users should provide the *dihedral* torsions that describe the most probable conformation because it is likely that the ring flip does not occur in AA-MD simulations, as shown in Figs. 4C and 4D.

### Lipid Backmapping

To test the performance of `mstool`, two CG lipid systems were transformed into AA ones. The first example was a low-resolution CG model in which four CG beads represent one lipid.^45,46^ Two types of lipids were involved in this example. One was a phospholipid, and another one was a neutral lipid triolein. I backmapped a CG membrane structure in which triolein nucleated inside a POPC bilayer (Fig. 5A). Although the resolution of the CG model was low, making a backmapping input file for this system was straightforward because `mstool` simply requires a mapping scheme and isomeric information of these two types of lipids. The conversion increased the resolution of the structure 34-fold without any bridging resolutions, and the resulting structure had no flipped isomers, demonstrating the robustness of `mstool`.

**Figure 5.**
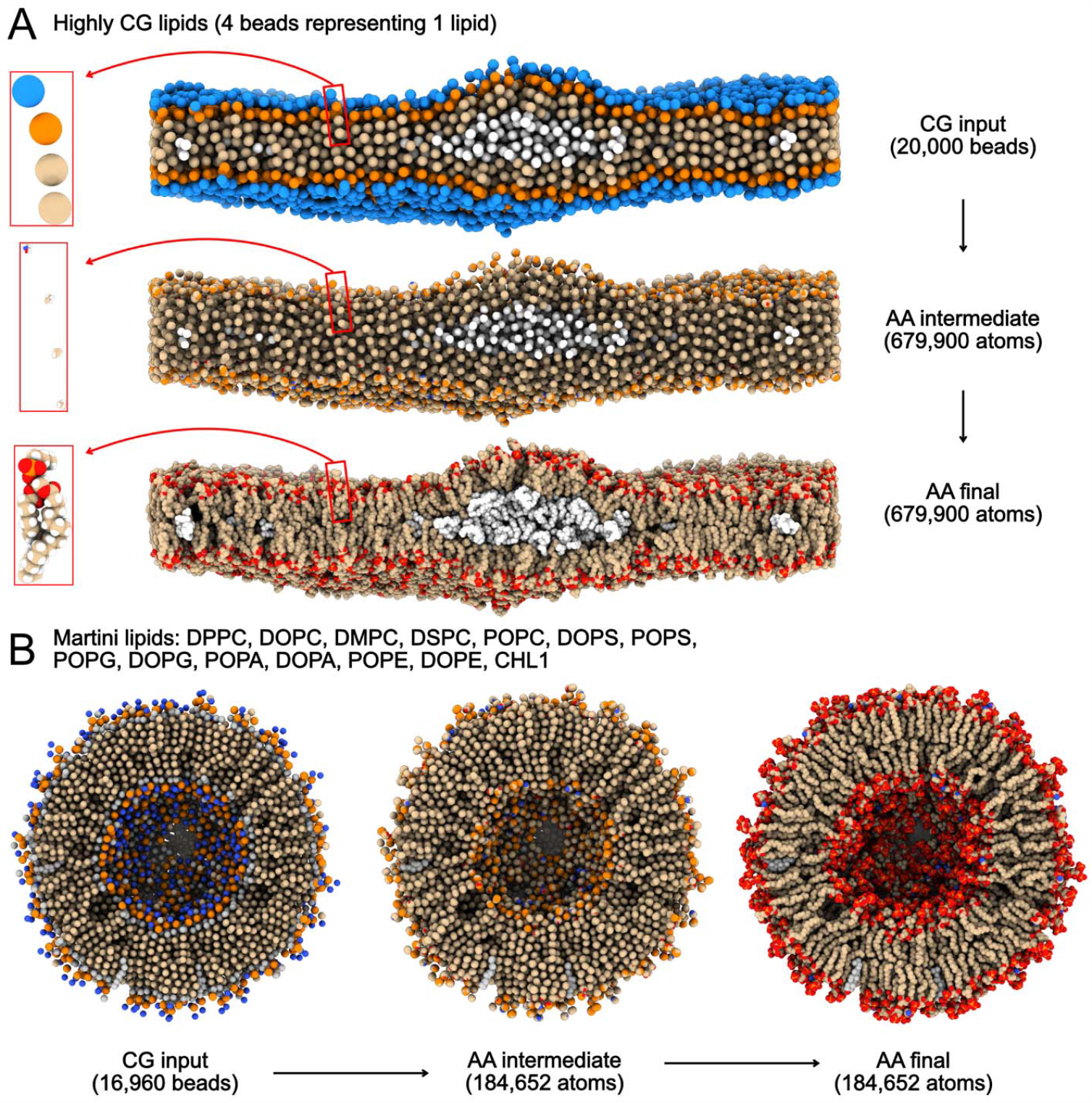
Lipid backmapping of (A) highly CG lipids and (B) Martini lipids.

Martini lipids are one of the most popular CG models.^18–21^ It has been shown that they accurately capture the physics and chemistry of many biological processes and reproduce the bilayer properties such as thickness and area per lipid.^47,48^ To test the efficacy of `mstool`, I backmapped a spherical bilayer at Martini resolution with 14 different Martini lipids into AA resolution (Fig. 5B). The backmapped structure had no flipped isomers. This shows the robustness of isomeric torsions applied during REM (Fig.1B and Fig. 1C) as each cholesterol molecule (residue name CHL1) has eight *chiral* centers, and a majority of lipids contain one or more *cis* bonds.

### Protein Backmapping

`mstool` supports multiple backmapping options for protein. The equilibration timescale of protein is much longer than that of lipids and usually beyond timescales attainable by AA-MD. Therefore, one should carefully choose a protein backmapping scheme that best fits the problem at hand. If there is an initial or reference AA protein structure, it is beneficial to incorporate it into the backmapping procedure. If there are no big conformational changes during CG-MD simulations, users can directly align the AA protein structure against the CG protein structure and then provide the aligned AA structure into `mstool` during REM. If there is a conformational change during CG-MD simulations, one can align only the rigid parts of protein and then build loops using a loop modeler provided in `mstool` or other software. However, random placement or geometrical projection can be an option if there is no initial AA protein structure, or the final CG structure is very different from the AA structure. Backmapping of T-cell intracellular antigen 1 (PDB: 2MJN) from Martini resolution to AA resolution was used as a test case.^49^ A short CG-MD simulation was performed to stray away from the initial AA structure (Fig. 6A).

**Figure 6.**
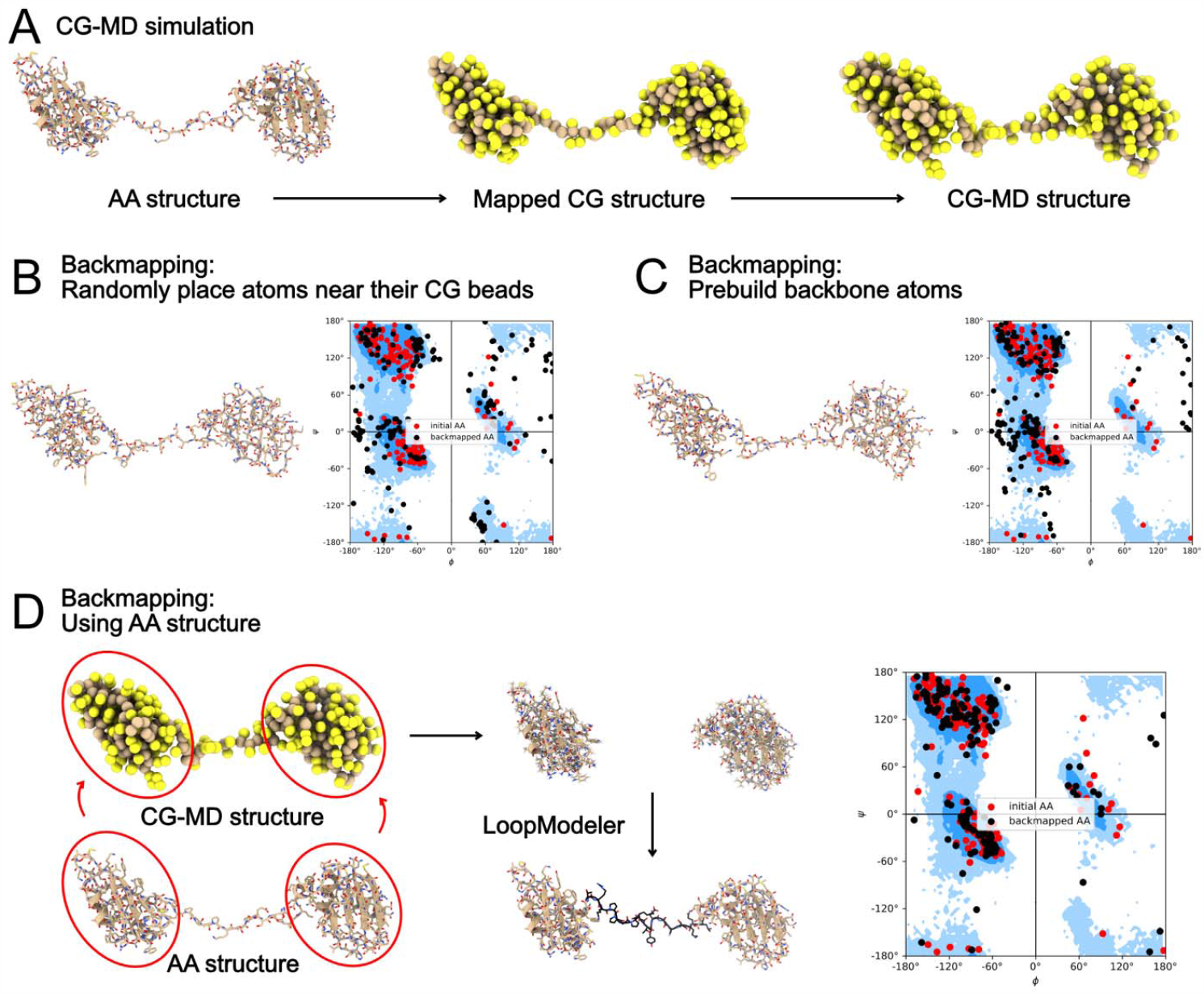
Protein backmapping. (A) CG-MD simulation of T-cell intracellular antigen 1. (B) First option. (C) Second option (default in `mstool.Ungroup`). (D) Third option using AA structure and `mstool.LoopModeler`.

The first option is to randomly place atoms of each amino acid at their corresponding CG beads, just like lipid backmapping, through `mstool.Ungroup`. For instance, the Martini resolution has one backbone bead (BB) per residue; Therefore, backbone atoms (N, HN, CA, HA, C, O) are initially placed at the location of their corresponding BB bead. Side chain atoms are also ungrouped similarly. An intermediate AA structure is relaxed (`mstool.REM`) and reviewed (`mstool.CheckStructure`).

The second option, the default option in `mstool`, is only slightly different from the first option in ungrouping. Instead of randomly placing AA atoms from CG beads, it predicts the positions of backbone atoms using the previously published geometrical algorithm (`CG2AA` and `Backward`).^22,23^ In short, it uses the positions of three consecutive backbone beads to obtain the positions of backbone atoms. Side chain atoms are placed near their corresponding CG beads. The rest of the procedure (`mstool.REM` and `mstool.CheckStructure`), is the same.

Backmapped protein structures using the first and second options are shown in Fig. 6B and Fig. 6C, respectively, along with their Ramachandran plots. Although both options provide a reasonable structure, there are some Ramachandra outliers. Therefore, I highly recommend the last option, which leverages the initial AA structure.

Most of cases, protein AA structures are present before running CG simulations. The last option of backmapping CG protein structures into biologically sound AA ones is to use these reference AA structures. While the Martini protein FF allows global changes in protein tertiary structures, it is not expected to change any secondary structures by design. If CG-MD simulations are run with an elastic network model (ENM), it is more obvious that protein structure changes little. Therefore, one can copy a rigid domain of an atomistic structure and align it with its corresponding CG counterpart. Loops that connect the rigid domains can be built using `mstool.LoopModeler` or other loop modelers. For instance, T-cell intracellular antigen 1 has two rigid domains, connected by a single loop. Therefore, it is reasonable to use the reference AA structure of the two domains in the backmapping process. The backmapped structure using this approach has fewer Ramachandran outliers (Fig. 6D).

### Membrane Protein and Lipid Backmapping

Backmapping a membrane protein and lipid system is not different from backmapping either lipid (Fig. 5) or protein (Fig. 6). However, it is important to note that protein and lipid should not be backmapped separately and then combined because a resulting structure will likely have unphysical contact between protein and lipid. All the atoms should be present during REM to get a final backmapped structure with no bad contacts. In this section, I used outer membrane protein F (PDB: 2OMF),^50^ facilitating the diffusion of molecules in the outer membrane of *E. coli*, as an example. A short CG-MD simulation was performed to stray away from an initial CG structure (Fig. 7A).

**Figure 7.**
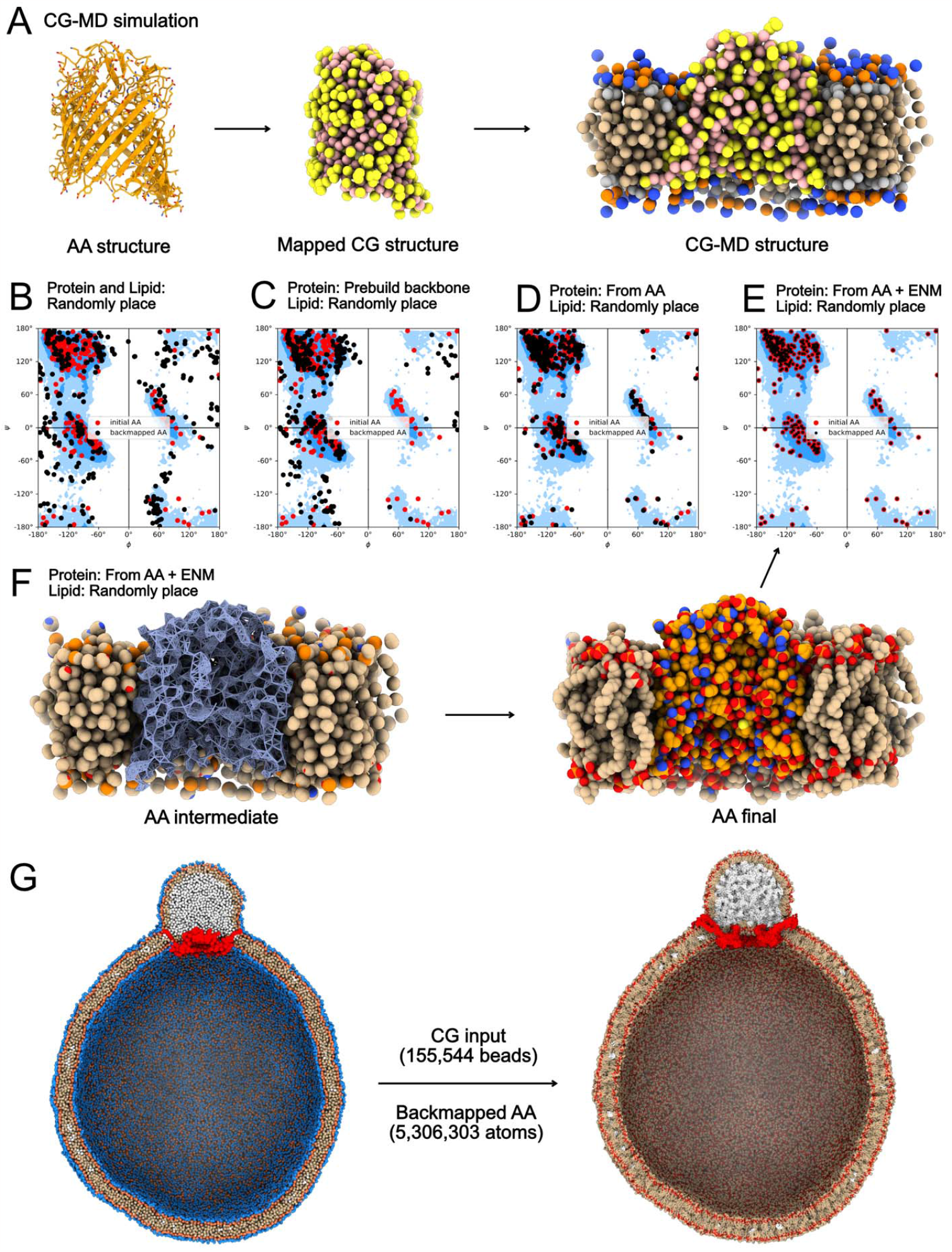
Lipid and protein backmapping. (A) CG-MD simulation of outer membrane protein F. (B-E) Ramachandran plots of the backmapped protein structures using various options. (F) Workflow that treats an input protein structure as a space-filling rock structure. (G) Heroic backmapping of lipid droplet biogenesis model. Red protein is seipin, and white is triolein.

The first approach randomly places protein and lipid atoms near their corresponding CG beads. The second option, the default, is the same as the first option except for protein backbone atoms, prebuilt based on three consecutive protein backbone beads in the ungrouping step. Both approaches gave a reasonable structure for the protein; However, as seen in the previous session, there were non-negligible Ramachandran outliers with these two options (Fig. 7B and Fig. 7C).

Considering the minimal conformational changes of the protein during the CG-MD simulation, it is reasonable to align the initial protein AA structure to the final CG structure and ungroup lipid AA atoms from lipid CG beads. However, because the initial AA protein structure is slightly different from the final CG protein structure, there are inevitably bad contacts between the initial AA protein atoms aligned to the final CG protein structure and lipid atoms ungrouped from CG lipid beads. Fortunately, REM is capable of relaxing such systems with unphysical contacts. The Ramachandran plot of a backmapped structure using the third option (Fig. 7D) showed fewer outliers compared to the first two options (Fig. 7B and Fig. 7C).

In the third option, protein topology is built by openMM during REM. If an input protein structure is not complete, has a modified protein residue such as glycosylation or phosphorylation, or has a ligand, openMM cannot create a protein topology. To deal with such cases, `mstool` has the last option to treat an input protein structure as a space-filler. In this case, the tool does not apply an AA protein FF to a provided protein structure but creates ENM for an input structure with moderate LJ parameters assigned to input protein atoms. During REM, non-protein atoms do not overlap protein atoms while protein maintains its position. Therefore, the Ramachandran plot of the final structure is identical to the initial AA protein structure (Fig. 7E); However, there are no bad contacts between protein and lipid atoms in a backmapped structure. The procedure of this option is illustrated in Fig. 7F.

`mstool` can work for larger and more complicated systems. Heroic backmapping of lipid droplet biogenesis is shown in Fig. 7G. It is a gigantic spherical bilayer with a diameter of 60 nm that shows seipin-mediated lipid droplet maturation.^51^ Protein is a seipin oligomer (PDB: 6DS5),^52^ facilitating triolein nucleation and proper lipid droplet maturation. This CG system is also a good example of protein backmapping using `mstool.LoopModeler`. The luminal domain of the seipin oligomer is rigid and changes little during CG-MD simulations. Therefore, I cut the luminal part of the AA protein structure and aligned it to the CG protein structure. In contrast, the transmembrane segments were flexible and became open up during CG-MD.^51^ Therefore, each transmembrane segment of the AA structure was aligned into its corresponding CG structure. A total of 22 loop structures that connect the luminal domain and the transmembrane segments were made using `mstool.LoopModeler` all at once. This way, while the AA structure of the luminal domain and transmembrane segments were preserved, the backmapped model also could capture the opening of seipin transmembrane segments during lipid droplet maturation.

## DISCUSSION

CG-MD simulations are helpful in understanding complex biological processes by allowing access to time and length scales beyond the scales typical of AA-MD simulations. However, CG models lack atomistic details critical in understanding protein structures, protein-lipid interactions, and protein-ligand interactions. Therefore, converting CG structures into AA ones in a biologically and physically sound manner is an important tool in the field of CG modeling of biophysical systems.

In this work, I have developed a backmapping tool that converts CG structures into AA ones and implemented it into a package called `mstool`. The procedure consists of three steps for resolution conversion. First, `mstool.Ungroup` simply places atoms near their corresponding beads. Second, `mstool.REM` applies a modified FF to a system and then performs energy minimization. The modified FF replaces the positive part of LJ with a cosine function and adds additional torsions to defined isomers. The former modification ensures numerical stability during energy minimization, and the latter does consistency in isomeric properties. Finally, a backmapped structure is reviewed by `mstool.CheckStructure`.

These three steps are decoupled. Therefore, each step can be combined with other backmapping tools. For instance, one can consider using approaches to create a better intermediate AA structure before energy minimization, such as fragment alignment^16,17^ or geometrical projection.^23^ Once an intermediate structure is obtained using either of these approaches or other methods, `mstool.REM` can relax the structure. Obviously, one can backmap CG structures into AA structures with other backmapping tools and then review flipped isomers with `mstool.CheckStructure`.

It should be noted that `mstool.REM` can be used for other purposes. For example, if an AA structure has multiple bad contacts, standard energy minimization will fail because of high potential energy. In such cases, `mstool.REM` can help resolve all the unphysical contacts so that standard energy minimization can be run afterward.

There are potential applications of this backmapping procedure that leverage the efficiency and fast equilibration of CG models. For example, `mstool.LoopModeler` can build a structure of missing loops without using structure templates. It is powerful to model any lengths of missing and multiple loops simultaneously. It is also simple to use, taking only a fasta file of protein chains and a protein structure file. The loop modeler automatically detects which parts are not present in the provided AA structure (except for the missing N-termini and C-termini) by comparing the sequence and residue number and builds loop structures. The details of the workflow and quality of loops will be discussed in a separate paper.

Another application is a membrane builder. CG membranes are easier to construct and equilibrate than AA membranes. One can build a CG membrane with a desired lipid composition at the Martini resolution, equilibrate the membrane at the CG level, and then backmap into the AA one. A prototype of a membrane builder is included in the repository.

## CONCLUSIONS

I present a robust and flexible backmapping tool with a minimal user input set: mapping and isomeric information. Leveraging the efficacy of the modified FF, the tool is capable of converting CG systems to AA ones.

## ACKNOWLEDGMENTS

I thank my Ph.D. advisor, Gregory A. Voth, for his mentorship and support. I am appreciative of my coworkers at D. E. Shaw Research for helpful discussion, Avi Robinson-Mosher, Peter Skopp, Robert McGibbon, Stefano Piana-Agostinetti, Qi Wang, Virginia Jiang, Dan Kozuch, Jim Valcourt, Parker de Waal, Jacob Pessin, and Brent Gregersen. Finally, I appreciate Won Hee Ryu, Gregory A. Voth, and anonymous reviewers for their time to review this paper.

## Notes

### Competing Interest Statement

The authors have declared no competing interest.

### Summary of Updates

Figure 1 revised; Figures 2-4 added; Comparison with other backmapping tools (CG2AT2 and Backward) added; Dihedral torsions introduced for ring conformation; Membrane equilibrium simulations added

